# Bacterial metabolic remodelling by convergent evolution in response to host niche-dependent nutrient availability

**DOI:** 10.1101/2024.09.04.611177

**Authors:** Amy C. Pickering, Jamie Gorzynski, Grace Taylor-Joyce, Willow Fox, Pedro Melo, Joana Alves, Hannah Schlauch, Fiona Sargison, Gonzalo Yebra, J. Ross Fitzgerald

**Affiliations:** The Roslin Institute and Edinburgh Infectious Diseases, University of Edinburgh, Easter Bush, Midlothian, Scotland, UK

## Abstract

New pathogens often arise after host jump events between species. However, our understanding of how bacterial pathogens pivot to distinct nutrient availabilities in a new host niche is limited. *Staphylococcus aureus* is a multi-host pathogen responsible for a global burden of disease in humans and farmed animals. Multiple human-to-bovine host switching events led to the emergence of *S. aureus* as a leading cause of intramammary infection in dairy cattle. Here, we employed *ex vivo* milk infections to investigate how bovine *S. aureus* has adapted to the dairy niche revealing metabolic remodelling including upregulation of genes for lactose utilisation and branched-chain amino acid biosynthesis in response to nutrient availability. Notably, infection of milk by bovine *S. aureus* results in a milk clotting phenotype associated with enhanced bacterial growth that is dependent on the protease aureolysin. The same adaptive phenotype has evolved convergently in different bovine *S. aureus* lineages via mutations in distinct regulatory gene loci that promote enhanced aureolysin expression. Taken together, we have dissected a key adaptive trait for a bacterial pathogen after a host-switch event, involving metabolic remodelling in response to the availability of nutrients. These findings highlight the remarkable evolutionary plasticity of *S. aureus* underpinning its multi-host species tropism.

## Introduction

The emergence of new pathogens often occurs after host-switching events between species, underpinned by adaptation to the new ecological niche. However, our understanding of the evolutionary processes and mechanisms driving bacterial host-adaptation remain limited. In particular, the capacity to adapt to distinct nutrient availability in the new niche is essential for bacterial proliferation and therefore critical for a successful host-switch [1]. Specialist bacteria have been reported to adapt through genome reduction to focus on particular nutrients available in a single niche [1–4]. In contrast, generalist bacteria may be required to maintain a broad range of metabolic pathways at their disposal for deployment simultaneously or sequentially [5]. There is increasing awareness of the complexities of nutrient availability for bacteria in different niches [6–8], and the role of nutritional immunity in limiting bacterial infection [9]. However, the ability of multi-host pathogens such as *Staphylococcus aureus* to pivot to distinct nutrients after a host switch is not well understood.

Human domestication of cattle for milk and associated dairy products provided a major niche for the expansion of bacteria, and evidence for dairy niche adaptation has been reported [10]. For example, *Lactococcus lactis,* used widely for production of cheese and buttermilk, exhibits enhanced lactose utilisation via plasmid-encoded lactose metabolism [11, 12], while the bovine intramammary pathogen *Streptococcus agalactiae* has acquired bovine-specific lactose and fructose operons [13, 14]. Furthermore, enhanced utilisation of casein, the major protein constituent of milk, is a feature of *L. lactis* [12] and mastitis isolates of *Escherichia coli* [15].

*S. aureus* is a leading cause of bovine mastitis in the global dairy industry and all contemporary lineages can be traced to a small number of independent host-switching events from humans which were followed by adaptation and clonal expansion [16, 17]. A combination of gene acquisition, loss and diversification have been reported to be associated with adaptation to the bovine niche including mobile genetic elements (MGE) encoding host-specific effectors of virulence and immune evasion [16, 18–20]. Bovine *S. aureus* strains also exhibit enhanced lactose utilisation compared to strains causing disease in other host-species [21, 22]. However, the functional evolutionary basis for *S. aureus* adaptation to the nutrients available in the dairy niche is poorly understood. Here, we have utilised a multi-omics approach to investigate the ability of *S. aureus* to successfully transition to a new ecological niche in the face of distinct nutrient availability. We discovered that bovine *S. aureus* has undergone metabolic remodelling since the host-jump, and that enhanced expression of the protease aureolysin is key to unlocking the nutrient potential of milk. This adaptive phenotype has evolved convergently in different *S. aureus* lineages via mutations in distinct gene loci, highlighting the remarkable capacity of *S. aureus* to expand into new host niches.

## Results

### Bovine *S. aureus* lineages exhibit a clotting phenotype associated with enhanced growth in milk

Previously, we identified that bovine *S. aureus* had increased capacity to utilise lactose (the major carbohydrate present in milk), in comparison to strains of human origin [22]. To investigate this observation further, we compared the growth phenotype of bovine and human *S. aureus* in bovine milk. Employing *ex vivo* milk infection experiments, we assessed 274 bovine and 91 human *S. aureus* strains from an array of common lineages (clonal complexes; CC) (Supp Table 1). Unexpectedly, 59.5% of bovine strains produced a clotting phenotype (separation into solid curds and liquid whey fractions) after 4-6 h incubation at 37°C, in comparison to only 4.4% of human strains (Fig 1A). Of note, the frequency of milk clotting varied in different bovine CCs with those that are globally distributed and long-established in ruminants e.g., CC97, CC133, and CC151 exhibiting highest clotting frequencies among bovine strains, of 76%, 84%, and 90%, respectively (Fig 1A). In comparison, those CCs more recently associated with cattle e.g., CC1, CC30, and CC188 tended to have lower frequencies of milk clotting among bovine strains with 40%, 25%, and 15%, respectively (Fig 1A) [16].

**Fig. 1.**
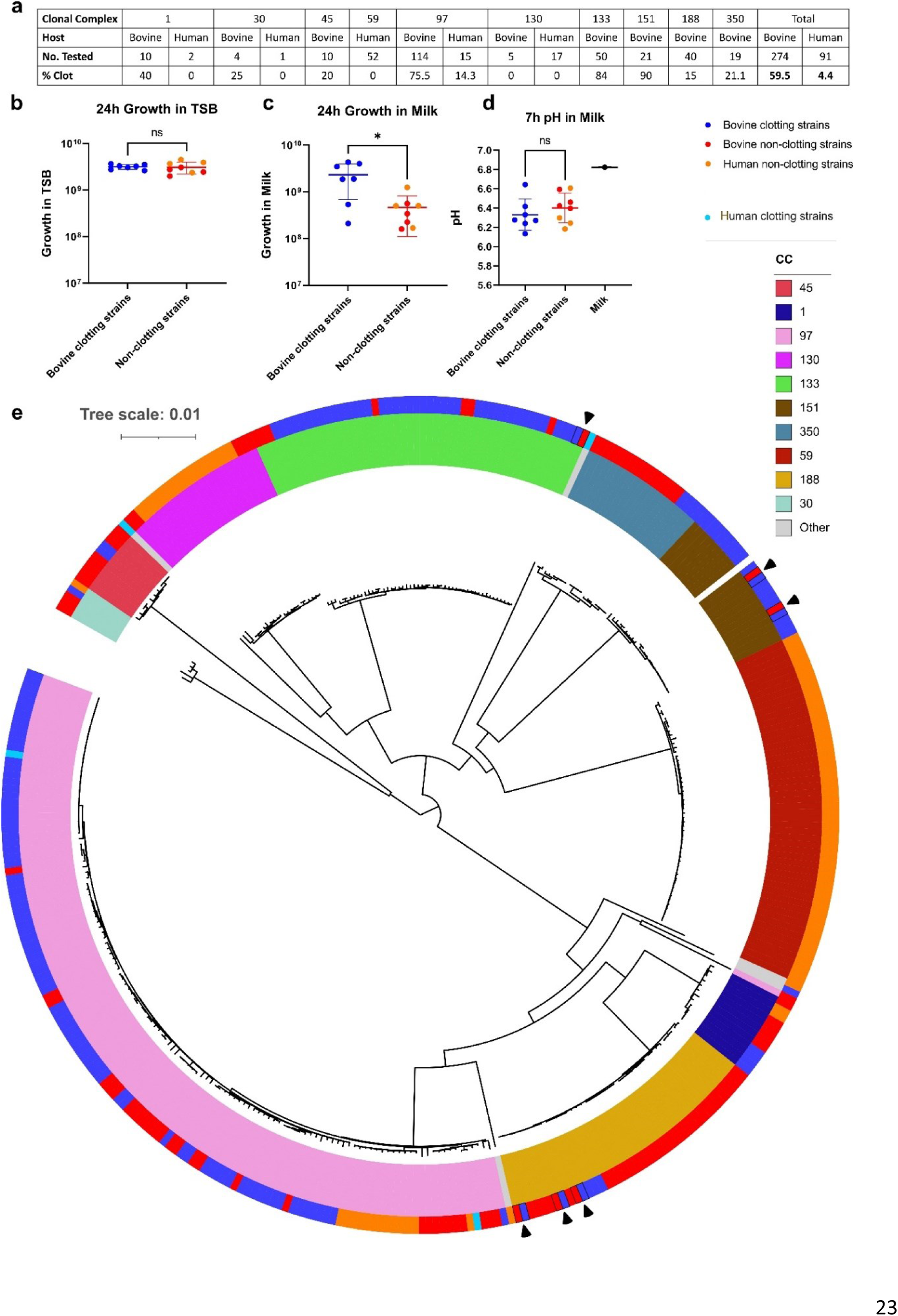
Bovine strains from multiple lineages have evolved a milk clotting phenotype associated with enhanced growth in milk. **a,** Frequency of the milk clotting phenotype in the isolates tested. **b&c,** CFU analysis of a diverse selection of bovine clotting, bovine non-clotting, and human non-clotting strains after 24 h growth in **b** TSB (two-sided Welch’s t-test, *t* = 0.2320, degrees of freedom = 9.838, *P* = 0.8213) and **c** milk (two-sided Welch’s t-test, *t* = 2.945, degrees of freedom = 6.482, *P* = 0.0235). **d,** pH analysis of the same strains after milk clotting was completed at 7 h post-inoculation. (two-sided t-test, *t* = 0.8601, degrees of freedom = 13, *P* = 0.4053). Asterisks indicate the level of significance: *P* > 0.05 (not significant, ns); 0.01 < *P* ≤0.05 (*). Each data point represents *n*=3 biological replicates for individual strains. Error bars, means ± standard deviation. **e,** A midpoint-rooted maximum-likelihood phylogeny of core SNPs across 365 isolates that were tested for clotting, corrected for ascertainment bias. Variants were called against a reference genome for the RF122 strain (AJ938182). Inner ring colours are used to indicate the clonal complex of isolates and outer ring colours indicate the milk clotting phenotype. Branches are drawn to the scale shown and measure the average number of substitutions per site. Black arrows are used to indicate pairs of isolates with contrasting phenotypes that were analysed in more detail, as referenced in the main text.

To investigate the potential relevance of the milk clotting phenotype for bacterial growth, seven selected bovine clotting strains (1xCC97, 3xCC133, 2xCC151, and 1xCC350), four bovine non- clotting strains (2xCC1, 1xCC97, 1xCC350) and four human non-clotting strains (2xCC97, 2xCC130) were cultured in both milk and TSB at 37°C. There were no differences in bacterial growth rate or yield of strains cultured in TSB (Fig 1B). However, bovine strains that clotted milk demonstrated an enhanced growth yield in milk compared to non-clotting strains of both bovine and human origin (Fig 1C). *S. aureus*-induced milk clotting is not due to milk acidification (curdling) as there was no significant reduction in pH of the milk media by the time clotting had occurred (Fig 1D). Overall, we have identified a novel milk clotting phenotype that was associated with enhanced growth, produced at high frequency by different global bovine *S. aureus* lineages.

### Phylogenetic analysis demonstrates a paraphyletic distribution of the milk clotting phenotype

To investigate the distribution and evolutionary origin of the milk clotting phenotype in *S. aureus*, we performed a maximum-likelihood phylogenetic analysis of genomes from 365 *S. aureus* strains from bovine and human origin, representing 10 lineages from 22 countries (Supp Table 1). The phylogeny was strongly delineated by CC, with most CCs demonstrating a paraphyletic distribution of the milk clotting phenotype (Fig 1E). For the three bovine lineages with the highest frequency of milk clotting (CC97, CC133, and CC151), the phylogeny indicates that the most recent common ancestor (MRCA) of each CC had the ability to clot milk, with sporadic loss-of-phenotype events occurring throughout the evolutionary history of each lineage (Fig. 1E). In contrast, for bovine lineages with lower frequencies of milk clotting (CC1, CC30, CC45, CC188 and CC350), the phylogeny indicates sporadic gain-of-phenotype events (Fig 1E). These data suggests that minimal genetic variations might be sufficient to confer or lose the milk clotting phenotype.

### Transcriptomic analysis of bovine *S. aureus* during growth in milk reveals adaptive metabolic remodelling

In order to investigate the basis for the enhanced growth phenotype in milk, we carried out RNA sequencing of 11 *S. aureus* strains from the CC97 global lineage including bovine clotting (n=4), bovine non-clotting (n=3) and human non-clotting strains (n=4) in the early (2 h) stages of growth in 50% milk/TSB media, in comparison to growth in TSB only. Strains were selected according to their distribution across the CC97 lineage, clotting and growth phenotype, and host-species origin (Supp Fig 1A-C).

RNAseq analysis was carried out as described in the Methods. Principal component and Kmeans clustering of global transcriptomic data revealed that bovine and human strains clustered together when cultured in nutrient-rich media (TSB) but formed two separate clusters when grown in milk; one cluster containing all bar one of the bovine strains with the second cluster containing all human strains and a single outlier bovine strain (Fig 2A). In TSB, only 51 genes were differentially expressed (p adjusted value of ≤0.05 and log2 fold change of ≤-1, ≥1) between experimental groups, with the majority more highly expressed in the bovine clotting strains compared to the human non-clotting strains including seven genes involved in lactose metabolism (*lacABCDFEG*), four genes involved in arginine biosynthesis (*argB*, *argC, argH* and *argJ*) and four genes involved in pyrimidine metabolism (*carB*, *pyrAA*, *pyrE* and *pyrF*) (Fig 2B-C; Supp Table 2). When cultured in milk, a total of 726 genes were differentially expressed (p adjusted value of ≤0.05 and log2 fold change of ≤-1, ≥1) between experimental groups (Fig 2D). We decided to focus on the 148 genes in milk that were differentially expressed with a log2 fold change of ≤-2, ≥2 for further analysis (Supp Table 3). Of the 89 genes annotated to KEGG pathways, 71.9% are predicted to have a role in metabolism. All bovine strains, irrespective of milk clotting phenotype, demonstrated elevated expression of valine, leucine and isoleucine (branched chain amino acids; BCAA) biosynthesis, arginine biosynthesis, and galactose metabolism in comparison to human strains (Fig 2E&G, Supp Fig 2B & Supp Fig 3B). Down-regulation of expression of urease and purine metabolism genes was observed in bovine clotting strains compared to non-clotting bovine and human strains (Fig 2E-F, Supp Fig 2 & Supp Fig 3) while bovine non-clotting strains exhibited increased phosphate transport expression (*pstSCAB*) associated with nucleotide biosynthesis (Fig 2F-G) [23]. Finally, we found that *S. aureus* clotting strains exhibited an elevated expression of genes encoding extracellular proteases in both milk and TSB in comparison to non-clotting strains from both cattle and humans (Supp Table 2 & 3). Overall, these data infer that bovine *S. aureus* has undergone adaptation to facilitate increased utilisation of lactose, to promote the synthesis of BCAAs and arginine, and increase the expression of extracellular proteases. We propose that the metabolic remodelling observed has contributed to enhanced growth of *S. aureus* in the dairy niche.

**Fig. 2.**
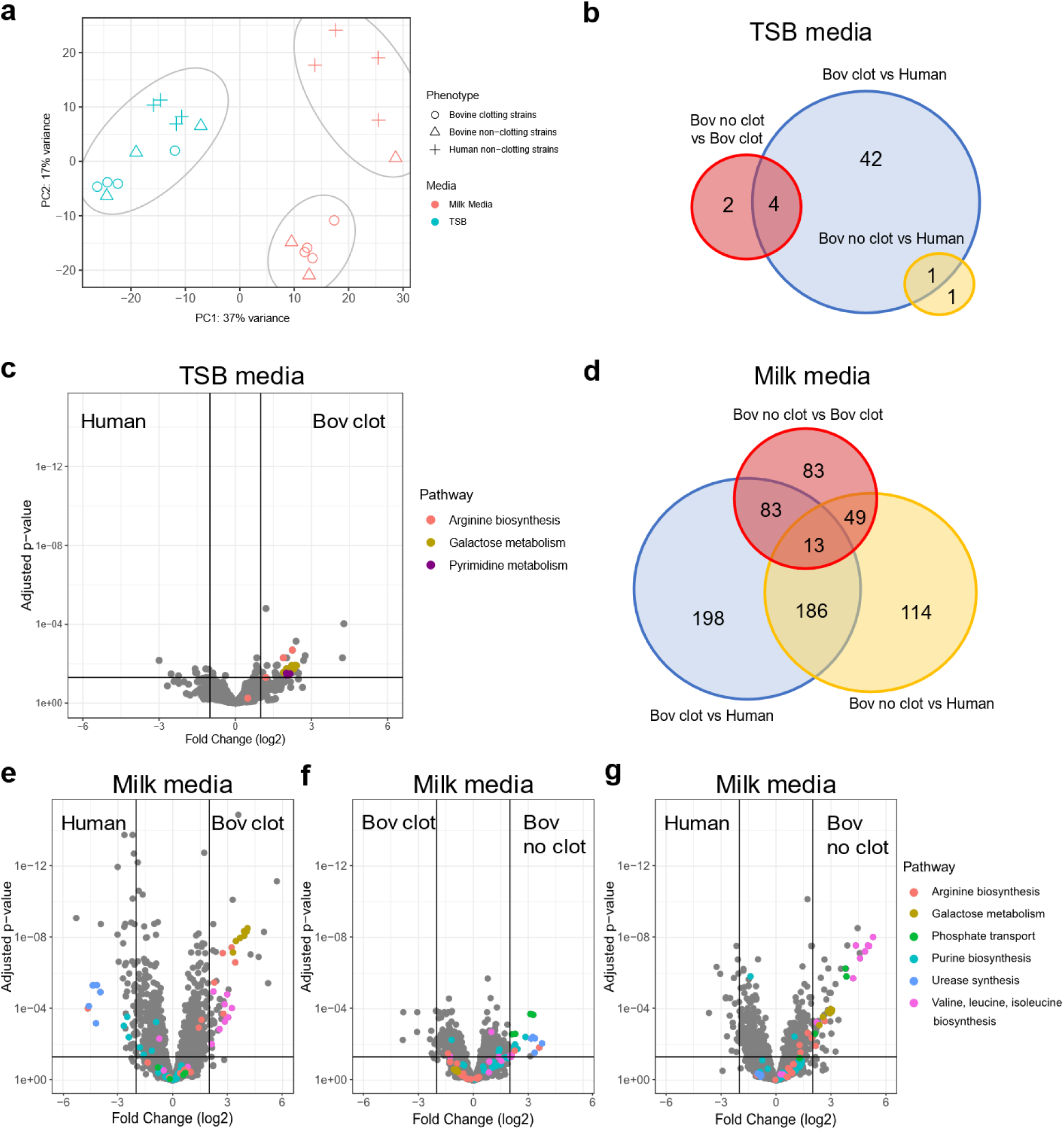
Transcriptomic analysis demonstrates metabolic differences between clotting and non-clotting strains in milk. **a,** Principal component and Kmeans clustering analysis, each separate cluster is circled in grey. **b,** Venn diagram showing differentially expressed genes in TSB. **c,** Volcano plot demonstrating the metabolic differences between bovine clotting and non-clotting strains in TSB. **d,** Venn diagram showing differentially expressed genes in 50% milk/TSB media between experimental groups. **e,** Volcano plot demonstrating the metabolic differences between bovine clotting and non-clotting strains in 50% milk/TSB. **f,** Volcano plot demonstrating the metabolic differences between bovine clotting and human non-clotting strains in 50% milk/TSB. **g,** Volcano plot demonstrating the metabolic differences between bovine non-clotting and human non-clotting strains in 50% milk/TSB. Differential expression was determined as P_adj_ of ≤0.05 and log2fold change of ≤-1, ≥1 for b-d and log2fold change of ≤-2, ≥2 for e-g.

**Fig. 3.**
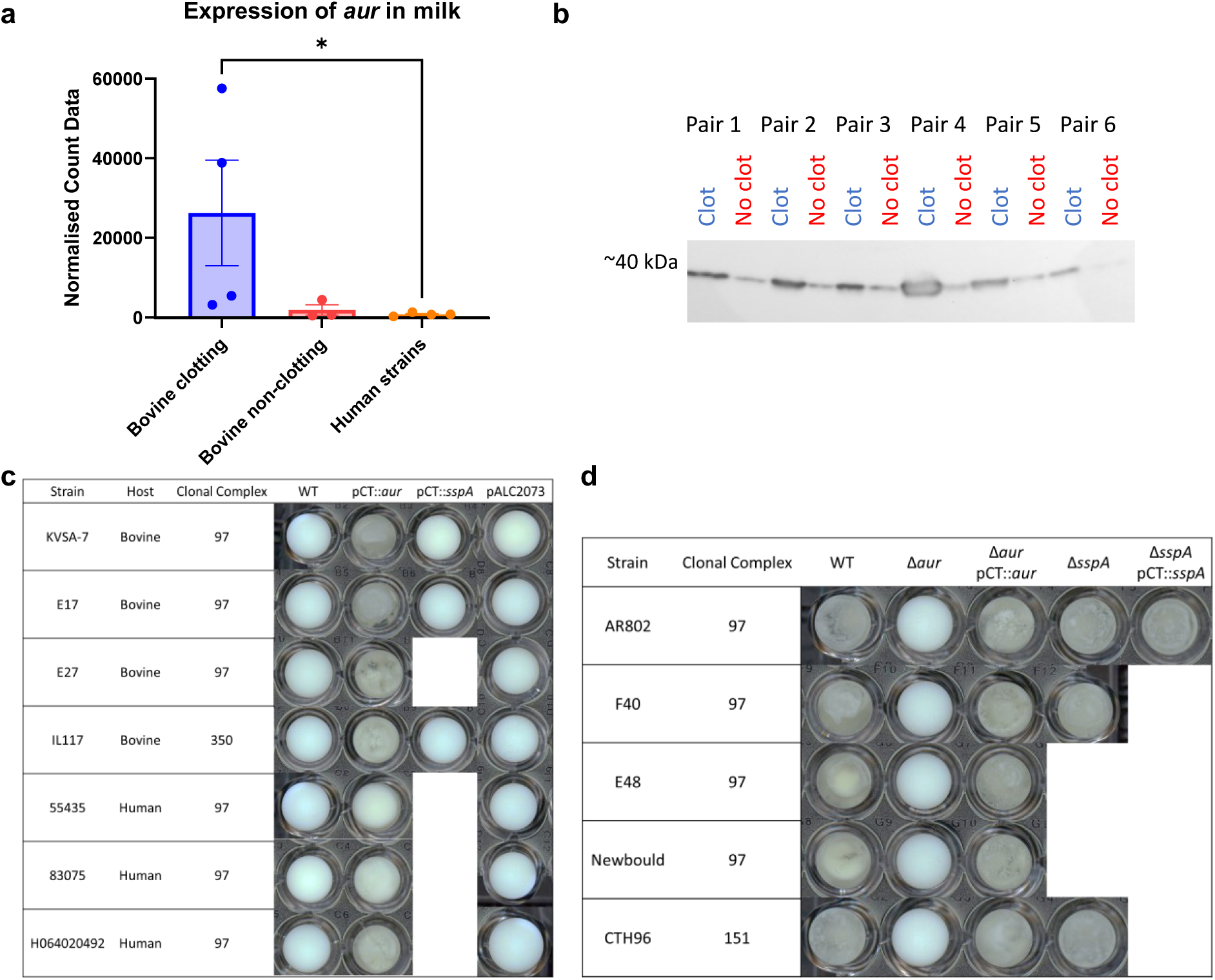
Increased expression of aureolysin in milk is required for milk clotting. **a,** Normalized count data of Aur expression after 2 h of incubation in 50% milk/TSB for each experimental group (Kruskal-Wallis statistic = 6.053, *P* = 0.0348). **b,** Western blot analysis of concentrated supernatant from paired isolates probed with anti-Aur IgY. **c,** Milk clotting phenotype of strains capable of clotting milk, deletion mutants and complemented strains after 24 h at 37°C under static conditions in a 96-well plate. **d,** Milk clotting phenotype of strains not capable of clotting milk, and complemented strains after 24 h at 37°C under static conditions in a 96-well plate. Each data point represents individual strains. Error bars, means ± SEM.

### The milk clotting phenotype is mediated by the protease aureolysin

Of note, the protease aureolysin (Aur) gene (*aur*), exhibited 14.5-fold increased expression in bovine clotting strains compared to bovine non-clotting strains, and 37-fold increased expression compared to human non-clotting strains during growth in milk (Fig 3A). The strong correlation between the clotting phenotype and *aur* gene expression led us to investigate Aur protein expression levels using bespoke Aur-specific antibodies (Eurogentec). From the phylogeny, six pairs of closely-related strains (<100 SNPs) with opposing milk clotting phenotypes were examined. Strikingly, for each pair, Aur protein expression was elevated in the clotting strain compared to the related non-clotting strain (Fig 3B).

To determine if this enhanced expression is responsible for milk clotting, we initially complemented a mutant derivative (USA300Δprotease [24]) of USA300 *S. aureus* strain (LAC) deficient in expression of all known extracellular proteases with the *aur* gene and its native promoter cloned into pCT (a derivative of pALC2073). USA300Δprotease pCT::*aur* mediated clotting of milk after 2 h of induction, whereas USA300Δprotease, USA300Δprotease pALC2073 (empty vector), and USA300Δprotease pCT::*sspA* (encoding the protease SspA) were unable to clot milk after 24 h (Supp Fig 4). Furthermore, 100 µg of concentrated supernatant from USA300Δprotease pCT::*aur* (Supp Fig 5) was sufficient to clot 5 ml of milk within 10 min. These data indicate that overexpression of Aur in the supernatant of an extracellular protease-deficient strain background is sufficient to mediate milk clotting. This was further examined through the overexpression of Aur via pCT::*aur* in six CC97 non-clotting strains of bovine origin, three of human origin and one CC350 strain, each of which was sufficient to promote milk clotting (Fig 3C).

**Fig. 4.**
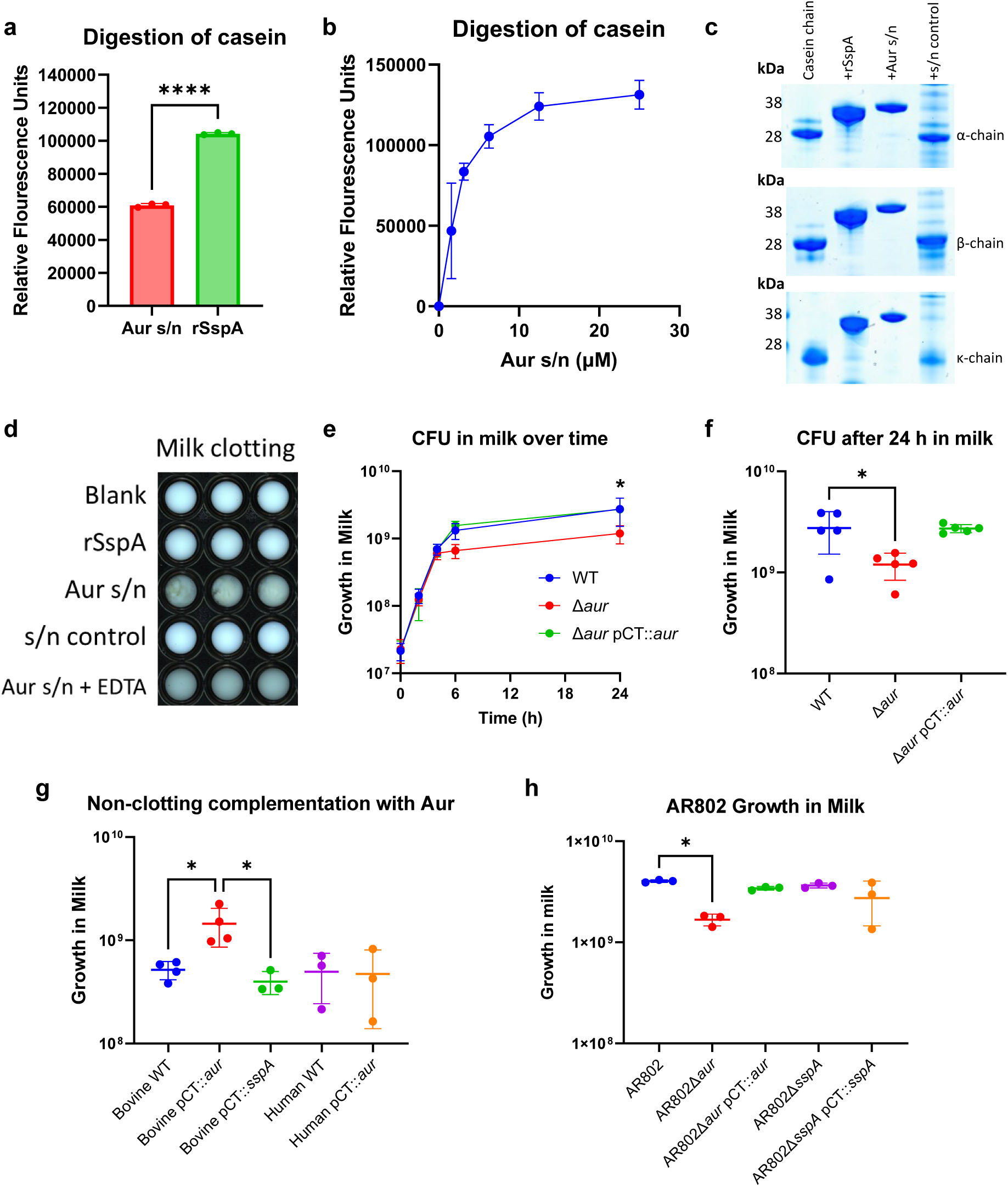
Aureolysin digests casein and promotes enhanced growth in milk. **a,** FTC-Casein digestion by 1 µM recombinant Aur, recombinant SspA, and Aur present in supernatant as measured by FRET (one-way ANOVA with Tukey’s multiple comparison test, *F* = 1281, *P* <0.001). **b,** Dose-dependent digestion of FTC-casein by Aur present in supernatant of USA300Δprotease pCT::*aur* as measured by FRET. Each data point represents *n*=3 replicates. Error bars, means ± standard deviation. **c,** digestion of individual chains of casein after 1h30 min incubation at room temperature as evaluated by SDS-PAGE. **d,** Milk clotting of 1 µM recombinant SspA, and Aur present in the supernatant after incubation at 37°C for 24 h under static conditions. 25mM EDTA was included as a negative control to inhibit Aur activity. **e,** CFU analysis over time in milk of bovine clotting strains, corresponding *aur* deletion mutants, and complemented Aur strains. **f,** Growth at 24 h of the same strains used in e (matched one-way ANOVA with Dunnet’s multiple comparison test, *F* = 11.42, adjusted *P* = 0.0293). **g,** CFU analysis of bovine and human non-clotting strains compared to those complemented with either Aur or SspA after 24h growth in milk. (one-way ANOVA with Tukey’s multiple comparison test, *F* = 6.076, *P* = 0.0065). **h,** CFU analysis of AR802 wild type, deletion mutant, and complemented strains after 24h growth in milk. (Kruskal-Wallis statistic = 16.53, approx. *P* = 0.0112). Each data point represents *n*=3 biological replicates for individual strains. Error bars, means ± standard deviation.

**Fig. 5.**
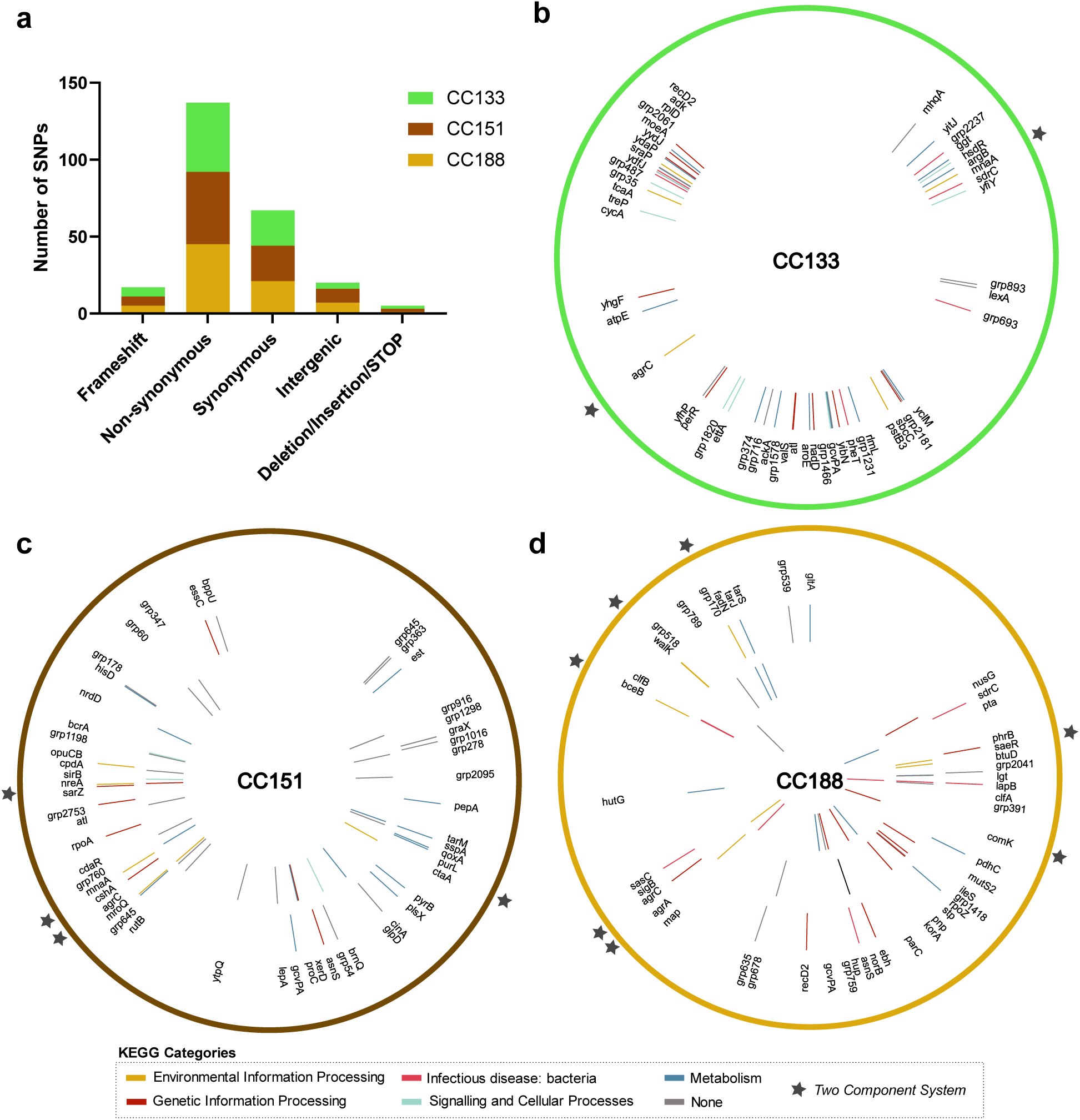
Distribution of SNPs among paired isolates and common pathways associated with milk clotting. **a,** The overall number of SNPs identified in the paired SNP analysis is provided in the context of each clonal complex examined. **b-d,** Non-synonymous (NS) SNPs of each pair are mapped to their corresponding location in the RF122 genome and demonstrate the spread of SNPs throughout the genome of **b,** one CC133 pair, **c,** two CC151 pairs – CH350 vs CH354 on the outer ring and IL83 vs CH381 on the inner ring, and **d,** three CC188 pairs – K130 vs K124 on the outer ring, K132 vs K125 in the middle and K163 vs K162 on the inner ring. Each SNP is coloured based on the assigned KEGG pathway. The stars highlight the genes categorised as being involved in two-component system signalling. Genome maps were generated using Circos in the Galaxy platform.

To determine if Aur is essential for the milk clotting phenotype, *aur* deletion mutant strains were generated in five CC97 bovine clotting strains, including the archetypal bovine strain Newbould (ST115), and the CC151 strains CTH96. Of the five Aur-deficient strains, four were unable to clot milk after 24 h at 37°C and the other was significantly delayed (Fig 3D). The clotting phenotype was restored by complementation with pCT::*aur* (Fig 3D). As Aur is known to trigger a proteolytic cascade and is required for the activation of SspA, which subsequently activates SspB [25], we considered that the ablation of the clotting phenotype in the *aur* deletion mutant could alternatively be explained by a lack of SspA activation by Aur. Accordingly, deletion mutants of *sspA* were generated in three strains (AR802 and F40 from CC97 and CTH96 from CC151). All three *sspA* mutants clotted milk in a manner similar to the wild type strains, confirming that Aur is essential for the milk clotting phenotype, and this is independent of SspA (Fig 3D). Finally, overexpression by complementation with pCT::*sspA* was not sufficient to induce milk clotting by any strain (Fig 3C-D). Taken together, these data indicate that enhanced Aur expression by *S. aureus* is sufficient and required to mediate the milk clotting phenotype and is independent of downstream proteases.

### Aureolysin mediates digestion of casein and is required for enhanced growth of bovine *S. aureus* strains in milk

Our transcriptomic and experimental data suggest that bovine strains have adapted to the ecological niche of the udder through metabolic remodelling and enhanced Aur expression. As casein digestion has been reported to promote milk clotting [26], we considered that Aur-dependent milk clotting may be mediated by casein digestion. To test this hypothesis, we examined the digestion of fluorescently-labelled casein using fluorescence resonance energy transfer (FRET) (Pierce^TM^). As SspA is known to digest casein [27] it was used as a positive control. The concentrated supernatant of USA300Δprotease pCT::*aur* (Supp Fig 5) mediated degradation of all three chains of casein in a dose-dependent manner with 1.7x lower activity than SspA (Fig 4A-C). However, only the concentrated supernatant of USA300Δprotease pCT::*aur* induced milk clotting whereas both rSspA and a supernatant control were unable to clot milk (Fig 4D).

Next, we tested if Aur is required for enhanced growth in milk. Comparison of bovine *S. aureus* clotting wild type, *Δaur* and pCT::*aur* complemented strains demonstrated that loss of Aur expression resulted in reduced growth after 24 h in milk (Fig 4E-F). Similarly, overexpression of Aur but not SspA in non-clotting bovine strains (3xCC97 and 1xCC350) enhanced growth in milk (Fig 4G). In contrast, for human non-clotting strains (3xCC97), overexpression of Aur mediated clotting but did not result in enhanced growth (Fig 4G). These data are consistent with our global transcriptomic analysis that revealed metabolic remodelling by bovine *S. aureus* compared to human strains, that facilitates the Aur-dependent enhanced growth observed in milk. Taken together, these data provide evidence for the functional adaptive evolution of bovine *S. aureus* to the dairy niche since the host-swich from humans.

### Enhanced aureolysin expression is driven by convergent evolution in bovine *S. aureus* lineages

Having established the requirement for increased Aur expression in milk clotting and the enhanced growth phenotype of *S. aureus* across multiple bovine-associated lineages, we wanted to examine the evolutionary genetic basis of this phenotype. Initially, we performed two genome wide association studies (GWAS) of clotting v non-clotting isolates, one using core SNPs as predictive factors and the second using the presence or absence of genes. In both analyses, no intragenic or intergenic SNPs or coding sequences (CDS) were significantly associated with the milk clotting phenotype (Supp Fig 6). These data suggest that there is not a single gene or mutation that is responsible for the phenotype in different lineages. Next, using a bespoke bioinformatic pipeline, we investigated the genetic differences including SNPs, indels and accessory regions between six closely-related pairs of isolates that differed in clotting phenotype in three distinct lineages (CC133, CC151 and CC188) (Supp Table 4). There was variation in the presence of MGE with one CC133 pair differing by the presence of a Sa6 phage and one of the CC188 pairs differing by the presence of an Sa3 phage. Furthermore, the 3 CC188 pairs contain insertion sequence elements, the precise genomic locations of which could not be confirmed using the short-read sequence data. A total of 307 SNPs were identified collectively between isolates within each pair, 17 SNPs resulting in frameshifts, 137 non-synonymous (NS) SNPs with at least one amino acid change, 67 synonymous SNPs, 20 intergenic SNPs within 100 bp of the CDS start codon, with the remaining five SNPs representing deletions, insertions, or stop codons being gained or lost (Fig 5A & Supp Table 4). One gene *gcvPA* encoding a glycine dehydrogenase involved in multiple metabolic pathways contained distinct SNPs in different lineages but overall, the variation differentiating each pair was in distinct gene loci distributed around the genome (Fig 5B-D). From the 92 coding sequences affected by at least one NS SNP that could be assigned to KEGG categories, 35 (38.0%) have a role in metabolism, 25 (27.2%) in genetic information processing, 11 (12.0%) in environmental information processing, and 10 (10.9%) in signalling and cellular processes. Notably, at the level of KEGG pathways, we identified 14 genes (15.2%) predicted to be involved with gene regulation including the two-component systems (TCS); *walK, saeR, agrA,* and *agrC,* (Fig 5B-D) in addition to transcription factors *abrB*, *perR*, and *sarZ*. Several of these systems including SarZ, Sae, Agr and Walk have already been reported to influence expression of Aur directly or indirectly [28]. Accordingly, we speculate that these mutations may underpin the difference in Aur expression levels identified in different pairs and we are currently investigating this hypothesis. Importantly, the distinct genetic variation associated with the clotting phenotype among the lineages examined, indicates convergent evolution has led to the milk-adaptive phenotypes.

## Discussion

As a generalist, *S. aureus* has the capacity to adapt to a wide array of host species and has undergone numerous host-switching events during its evolutionary history, particularly between humans and cattle [22]. In total, 14 major contemporary bovine-associated lineages have been identified that have resulted from at least four human-to-bovine host jumps [29]. In some cases, host jumps back into humans have led to the emergence and spread of human epidemic clones [30]. Accordingly, *S. aureus* represents an excellent model for exploring the adaptation of a generalist bacterial species to a new host niche. Here we have identified that bovine strains of *S. aureus* have undergone adaptive metabolic remodelling compared to human strains of the same lineage (Fig 2). Our findings support a model whereby bovine *S. aureus* have evolved increased capacity to utilise the abundant carbohydrate lactose and synthesise BCAAs and arginine. Considering that free BCAAs and arginine are present in milk [31], it is unclear why these biosynthetic pathways should be upregulated in bovine compared to human strains, especially as they are not usually activated during *in vitro* conditions [32, 33]. Consistent with our findings, previous studies have reported elevated levels of BCAAs in milk during infection by *S. aureus*, and this is currently being explored as an alternative marker for subclinical mastitis [34]. In addition to protein synthesis, an array of important processes have been linked to BCAA biosynthesis, including the production of branched-chain fatty acids and the regulation of virulence by CodY [35] or Sae [36].

After milk clotting, underpinned by increased expression of Aur by bovine strains, the metabolic remodelling of bovine *S. aureus* promotes enhanced growth in milk (Fig 1C and 4E). The discovery of a novel Aur-dependent growth effect provides new understanding of how Aur contributes to nutrient acquisition in different ecological niches. Aur is already known to be important during abscess infection through the generation of collagen peptides as nutrients for growth, highlighting the broad relevance of Aur for the metabolic adaptation of *S. aureus* in different tissue types and host-species [37]. The importance of peptide metabolism by a human *S. aureus* strain in milk has been reported previously, with a second phase of growth attributed to the release of amino acids by casein digestion in an SspA- and oligopeptide transporter (Opp3)- dependent manner [38]. In the current study, we demonstrate that the enhanced growth in milk by bovine strains is Aur- rather than SspA-dependent (Fig 4F) and likely due to the proteolysis of casein (Fig 4A-C), the most abundant protein in milk. The elevated expression of Aur in milk clotting strains (Fig 3B) is not dependent on genetic alterations to the *aur* gene or its promoter but instead different mutations at distinct regulatory gene loci in different strain backgrounds have led convergently to the same phenotype. The regulation of Aur by *S. aureus* is highly complex with numerous integrated regulatory networks including the regulons of TCS, influencing Aur expression [28]. In the current study, SNPs in genes of the *agr* operon, encoding a quorum sensing transcriptional regulator that influences the expression of a wide array of virulence factors including proteases [39], were present in four of the six paired isolates. For example, truncations in *agrC* were identified that are predicted to cause *agr* dysfunction in the non-clotting isolates of the two CC151 pairs. The mutations of *agr* or the other affected TCS may be responsible for the differential expression of Aur in bovine strains with differing milk clotting phenotypes. However, Aur gene expression could also be impacted by the presence of phage or the transposable elements in differing gene loci, and our data support numerous possible evolutionary trajectories to the same adaptive phenotype.

Although, the great majority of bovine strains from different lineages had the clotting and growth enhancement phenotype, the dominant established global clones CC97 and CC151 had a higher frequency than the more recently emerged and more locally distributed clones e.g., CC1, CC30, CC45, CC188 and CC350. The adaptation of these clones to the dairy ecological niche may still be ongoing [16], or it is also feasible that increasing protein translation and secretion associated with the clotting phenotype may have a fitness cost in non-milk-associated niches during bovine skin colonisation or intracellular invasion, thereby selecting for sporadic loss of function.

In conclusion, our data reveal how bovine *S. aureus* has undergone convergent remodelling of existing metabolic pathways to pivot to distinct nutrient availabilities in the bovine host. These findings highlight the evolutionary plasticity of *S. aureus* that promotes its remarkable versatility and capacity to expand into new niches.

## Methods

### Bacterial strains

All isolates employed for genetic and phenotypic analysis are listed in Supp Table 1. Mutant strains generated in this study are described in Supp Table 5. *S. aureus* strains were routinely cultured in Tryptone Soya Broth (TSB) at 37°C with shaking at 180 rpm or statically on Tryptone Soya Agar (TSA). Supplementation with 10 µg/ml chloramphenicol was used as appropriate. For cloning, *Escherichia coli* strains were cultured in Luria Broth (LB) supplemented with 100 µg/ml ampicillin or 15 µg/ml chloramphenicol.

### Milk clotting phenotype and assessment of pH

Arla Cravendale filtered whole milk was inoculated with 1:100 TSB overnight culture, concentrated supernatant or recombinant proteins and incubated for up to 24 h at 37°C; 180 rpm. The pH was measured by centrifuging the sample and using a pH meter to measure the pH of the whey or unseparated milk. A clotting phenotype was assigned if separation of the milk into whey and clot was observed or if a solid milk clot had been produced upon tilting the tube. Each strain was tested in three different milk sources.

### Growth analysis in milk

Overnight TSB cultures were standardised to a starting OD_600_ of 0.05 in 500 µl of Arla Cravendale filtered whole milk and incubated at 37°C with shaking at 180 rpm for the noted time. For some experiments, 500 µl TSB was inoculated in the same manner as a comparison. At each time point, the presence of milk clots were noted before addition of 0.02 % (w/v) SDS and 10 min vortex at max speed to separate any clots before CFU analysis. Each strain was tested in three different milk batches.

### Data collection

For genomic analysis, we included 358 whole genome sequence (WGS) datasets that were previously uploaded to public repositories under the codes listed in Supp Table 1. Seven additional WGS datasets were uploaded to NCBI under the new project code PRJNA1153292. For 347 sets of reads, we used the draft genome assemblies that were generated previously [16, 22, 40], which were assembled from reads following the protocols described in their respective projects as indicated in Supp Table 1. The remaining 18 read sets were trimmed using Trimmomatic (v0.36) and assembled using SPAdes (v3.13.0). Prior to variant calling, we checked the assembly quality of all collected genomes with QUAST (v4.6.3) [41] using the RF122 reference genome (AJ938182) to measure assembly statistics.

### Phylogenetic analysis

We used snippy (v4.4.5, https://github.com/tseemann/snippy) to identify SNPs with a reference genome for RF122 (AJ938182), generating pseudo-reads from the draft assemblies with the --ctgs parameter. Using snippy-core to generate an alignment, we filtered out recombined regions from the alignment using gubbins (v2.3.4) [42] and extracted core SNPs using snp-sites (v2.5.1) [43]. The core SNPs were used to generate a maximum-likelihood phylogeny with IQtree (v2.0.5) [44], using a General Time Reversible (GTR) model of evolution with a four-site gamma distribution to model differences in the evolutionary rate between branches. The tree was generated using the ultra-fast bootstrap method with 5000 replicates and optimised on the bootstrap alignment using nearest neighbour interchange with the --bnni parameter. Ascertainment bias correction was applied to prevent the overestimation of branch lengths by excluding invariant sites and polytomies were collapsed with the --polytomy parameter.

### RNA sequencing experimental set-up and analysis

To facilitate RNA extraction, 50% Arla Cravendale filtered whole milk was used diluted in TSB. Overnight cultures were standardised to OD_600_ 0.1 in TSB cultures and OD_600_ 0.25 in 50% milk/TSB cultures in 5 ml volumes, incubated at 37°C at 180 rpm for 2 h before pelleting at 4863 x g for 5 min. The pellet was re-suspended in 200 µl TE buffer and transferred into a 96-well plate before centrifugation. The supernatant was removed and the pellets immediately frozen on dry ice containing 100% ethanol. The frozen pellets were briefly stored at -80°C before shipment to Genewiz, UK for RNA extraction and paired-end RNA sequencing.

Raw data reads have been deposited in ArrayExpress (E-MTAB-14384). Bowtie2 v2.4.2 [45] was used to map sequence reads to a corresponding indexed reference genome built using bowtie2- build. Before reads were mapped, Cutadapt v1.16 [46] trimmed adapters. Samtools v1.10 [47] was first used to convert the sequence alignment and map (SAM) files, outputted by Biowtie2, to a binary alignment and map (BAM) file, then to sort BAM files by genome position. CoverageBED (BEDtools v2.30.0) [48] generated count data files from sorted BAM files. Analysis of differential gene expression between groups of different strains, required shared genes across all genomes to be identified and count data files adjusted so that all shared genes had the same name across all count data files. To identify shared genes across all genomes, Roary v3.6.2 [49] was used to calculate a pan genome. The core genes were defined as genes with >95% sequence identity. For duplicated genes, Roary assigned multiple genes within a genome to a single core gene and the first gene listed by Roary was arbitrarily taken as the gene representing this core gene. Hence, only the count data for this gene, and not the other duplicates, were included in later analysis. In the later analysis of differential expression data, none of the genes of interest represented duplicated genes. The Deseq2 (Bioconductor v3.13) [48] package was used to generate differential expression data.

### Aureolysin Western blot analysis

Aur-specific IgY antibody was generated from immunization of two hens with two 16 amino acid peptides (CYYKDTFGRESYDNQG and CEGDALRSMSNPEQFG) of Aur by Eurogentec. Antibody was affinity-purified from eggs after three antigen boosts. For Western blot analysis, *S. aureus* was cultured for 5 h in TSB before centrifugation and concentration of supernatant using Amicon Ultra Centrifugal units (10 kDa cut-off). The Trans-Blot Turbo Transfer system (Bio-Rad) was used. Before blocking, Revert^TM^ 700 Total Protein Stain (LICOR) was used as a normalization strategy to allow quantification of Aur expression between paired isolates using the Licor Odyssey^®^ M. Blots were then blocked in 8% (w/v) semi-skimmed milk powder in PBS for at least 2 h at RT (Sigma Aldrich) before adding 15 µg anti-Aur IgY antibody overnight at 4°C. After washing, 2 µg F(ab) goat anti-chicken-IgG-HRP (Sigma) was applied for 1 h in PBS-Tween-20 0.05%. Reactive bands were visualised using ECL and the Syngene GeneGnome.

### Generation of protease expression and deletion mutant constructs

For protease expression constructs, *aur* or *sspA* genes were amplified with their native promoter and ribosomal binding site (Supp Table 6) using Q5 polymerase (NEB). Constructs were generated in the expression plasmid pCT digested with SacI and Eco*R*I and Antarctic phosphatase treated (NEB) using Gibson assembly (NEB), introduced in *E. coli* IM08B [50] and subsequently into relevant *S. aureus* strains (Supp Table 5). Constructs were confirmed by Sanger sequencing (Eurofins).

Allele replacement was performed using thermosensitive pIMAY-Z [50] using protocols previously described [51]. Briefly, oligonucleotide primers were used to amplify ∼500 bp flanking regions of *aur* or *sspA* genes (Supp Table 6) using Q5 polymerase (NEB) and *S. aureus* gDNA (Monarch genomic DNA purification kit, NEB). Gibson assembly (NEB) and transformation into *E. coli* DC10B [51] allowed insertion of AB and CD PCR products into pIMAY-Z digested with KpnI and NotI. After confirmation of construct generation by Sanger sequencing (Eurofins), plasmids were transformed into relevant *S. aureus* strains (Supp Table 5). After temperature shifting, gene deletions and the absence of spurious mutations that could impact phenotype were confirmed through whole genome sequencing (MicrobesNG).

### Digestion of casein

To generate Aur concentrated supernatant, USA300Δprotease pCT::*aur* was induced at mid-exponential phase with 3 µg/ml anhydrotetracycline and cultured overnight. Supernatant was concentrated using Amicon Ultra centrifugal units (10 kDa cut-off) (Supp Fig 5) and dialysed into PBS using a Float-A-Lyzer G2 (Spectrum Labs). Protein concentration was quantified using the DS-7 UV-Vis Spectrophotometer (DeNovix). The proteolytic activity of 1 µM of concentrated supernatant containing Aur was compared to recombinant SspA protein (Sigma) using FTC-Casein (Pierce Fluorescent Protease Assay Kit, Thermo Scientific). The manufacturer’s instructions for a microplate assay were followed with a 30 min reaction time before reading on the CLARIOstar plate reader (BMG LABTECH).

To assess the ability to digest all three casein chains, 2 µg of each α-casein, β-casein, and κ-casein chains of bovine origin (Sigma-Aldrich) were incubated with 4 µg of protein for 1h30 min at room temp and then analysed by SDS-PAGE (4-20% resolving gel, Bio-Rad) for cleavage. 25 mM EDTA was used as a control to inhibit the activity of Aur.

### Genome-wide association analysis

GWAS was performed using a linear mixed-effects model in pyseer (v1.3.6) [52] with the SNPs from the core SNP alignment as features. To control for population structure, a pairwise-kinship matrix was generated from the phylogeny using the phylogeny_distance.py pyseer script and provided as co-variates in the model using the --similarity parameter. A separate GWAS was performed by the same method for COGs that were calculated using panaroo (v1.1.2) [53]. For this GWAS, genomes were annotated using PROKKA (v1.14.6) [54] with the --compliant and -- force parameters, the --centre parameter listed as ‘UoE’ and the --genus specified as “Staphylococcus”. The pangenome was then constructed from .gff files using a core threshold of 100% and the --alignment parameter set to ‘pan’ with other parameters left as default (a gene identity threshold of 98%). To determine the significance associations in either GWAS, a 0.05 significance threshold was selected with a Bonferroni correction for multiple testing applied. Manhattan plots were generated using ggplot2 (v3.4.4) in RStudio (v4.4.2).

### Detecting paired genomic differences

To identify genomic differences between pairs of isolates, variant calling was performed with snippy (snippy v4.4.5), using the PROKKA .gbk file for the non-clotting strain as the reference and the contigs for the clotting strain as the query with the --ctgs parameter. The closest gene was identified for each SNP using the closest script in bedtools (v2.29.2) [55], converting gene gff files to .bed files using AGAT (v0.8.0, https://github.com/NBISweden/AGAT) [56]. All gene sequences were extracted from PROKKA output using an in-house gff parsing script (https://github.com/JPegorino/gff-parser) and translated to protein sequence using the transeq script in EMBOSS (v6.6.0) [57]. KEGG pathways were then annotated from the NCTC8325 strain using the corresponding online BRITE database (https://www.genome.jp/brite/sao00001). To achieve this, the gene nucleotide FASTAs were matched with CDSs in NCTC8325 (ASM1342v1, downloaded from NCBI) using BLAST (v2.15.0), filtering by 98% identity and coverage. Each SNP location was mapped to the RF122 genome (AJ938182) and visualised using Circos in the Galaxy platform [58].

To identify larger indels and accessory regions, we used Mauve (snapshot 2015-02013) [59] to reorder the contigs in both isolates to best match the sequence in a closely related reference, selected from a database of complete *S. aureus* genomes in Refseq by minimum mash distance (calculated with mash v2.3 [60]). Links to each selected reference in Refseq are included in Supp Table 4. Aligned contigs were then combined into single sequences and compared manually using the Artemis Comparison Tool (Release 18.1.0) [61], creating .gff files that marked on the contig breaks by adapting the code in https://github.com/widdowquinn/scripts/blob/master/bioinformatics/stitch_six_frame_stops.py. To identify accessory genes, panaroo [53] was used for each genome pair and matched up the data to CDSs in the combined assembly FASTAs by re-annotating with PROKKA [54], keeping the same parameters except for providing the original annotations as trusted annotations for the -- proteins parameter.

## Acknowledgements

J.R.F. was funded by a response mode grant (BB/W014920/1) and an institute strategic grant (BBS/E/D/20002173) from the Biotechnology and Biological Sciences Research Council (United Kingdom), in addition to a grant from the Wellcome Trust (201531/Z/16/Z). The funders had no role in study design, data collection and interpretation, or the decision to submit the work for publication. We are grateful to Dr. Dara O’Halloran and Prof. Joan Geoghegan for providing the expression plasmid pCT.

## Supplemental Material

TABLE S1. The *S. aureus* strains tested for the milk clotting phenotype and used for phylogenetic analysis.

TABLE S2. Genes differentially expressed in TSB

TABLE S3. Genes differentially expressed in milk media

TABLE S4. Pairwise SNP analysis.

TABLE S5. Constructs used in this study. TABLE S6. Primers used in this study.

FIG S1. Milk clotting and growth of ST97 strains selected for transcriptomic analysis.

FIG S2. Volcano plots demonstrating the metabolic pathways differentially expressed between groups in milk.

FIG S3. Schematics of the major metabolic pathways differentially expressed between groups in milk

FIG S4. Overexpression of aureolysin provides milk clotting of *S. aureus* USA300

FIG S5. Concentrated supernatant of USA300Δprotease pCT::*aur*

FIG S6. Genome wide association analysis does not identify a genetic basis for the milk clotting phenotype within lineages, suggesting convergent evolution.

