## Supplemental Figures for "Bacterial metabolic remodelling by convergent evolution in response to host niche-dependent nutrient availability"

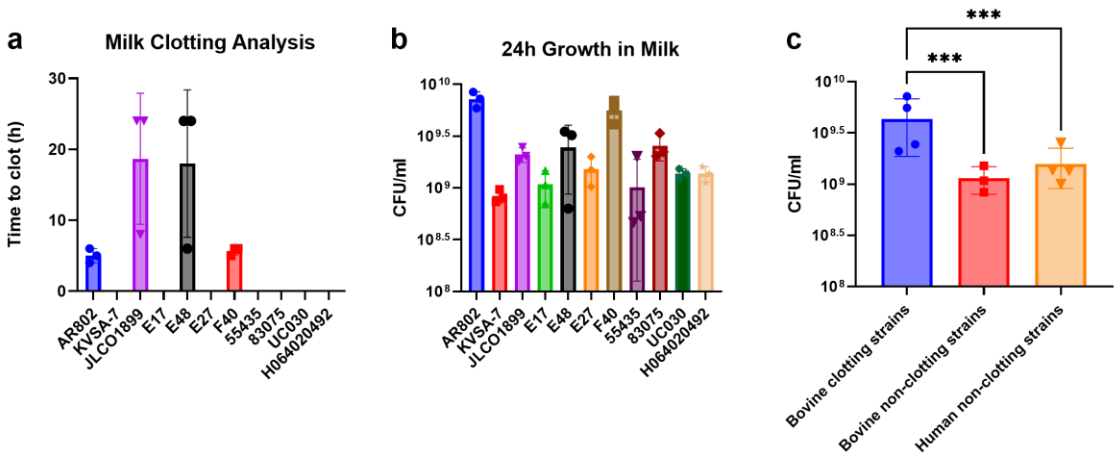


**Supp. Fig. 1| Milk Clotting and Growth of ST97 strains selected for transcriptomic analysis. a,** Time to clot milk of each strain. If no data points are present then the strain did not clot milk after 24 h of growth at 37°C with shaking. **b,** CFU analysis of strains cultured in 5 ml of Arla Cravendale filtered whole milk for 24 h at 37°C with shaking. Each data point represents a biological replicate, *n*=3. **c,** Data is combined for each milk clotting phenotype. Each data points represents *n*=3 biological replicates for each strain. Error bars, means ± standard deviation. Two-way ANOVA, *** p<0.005.

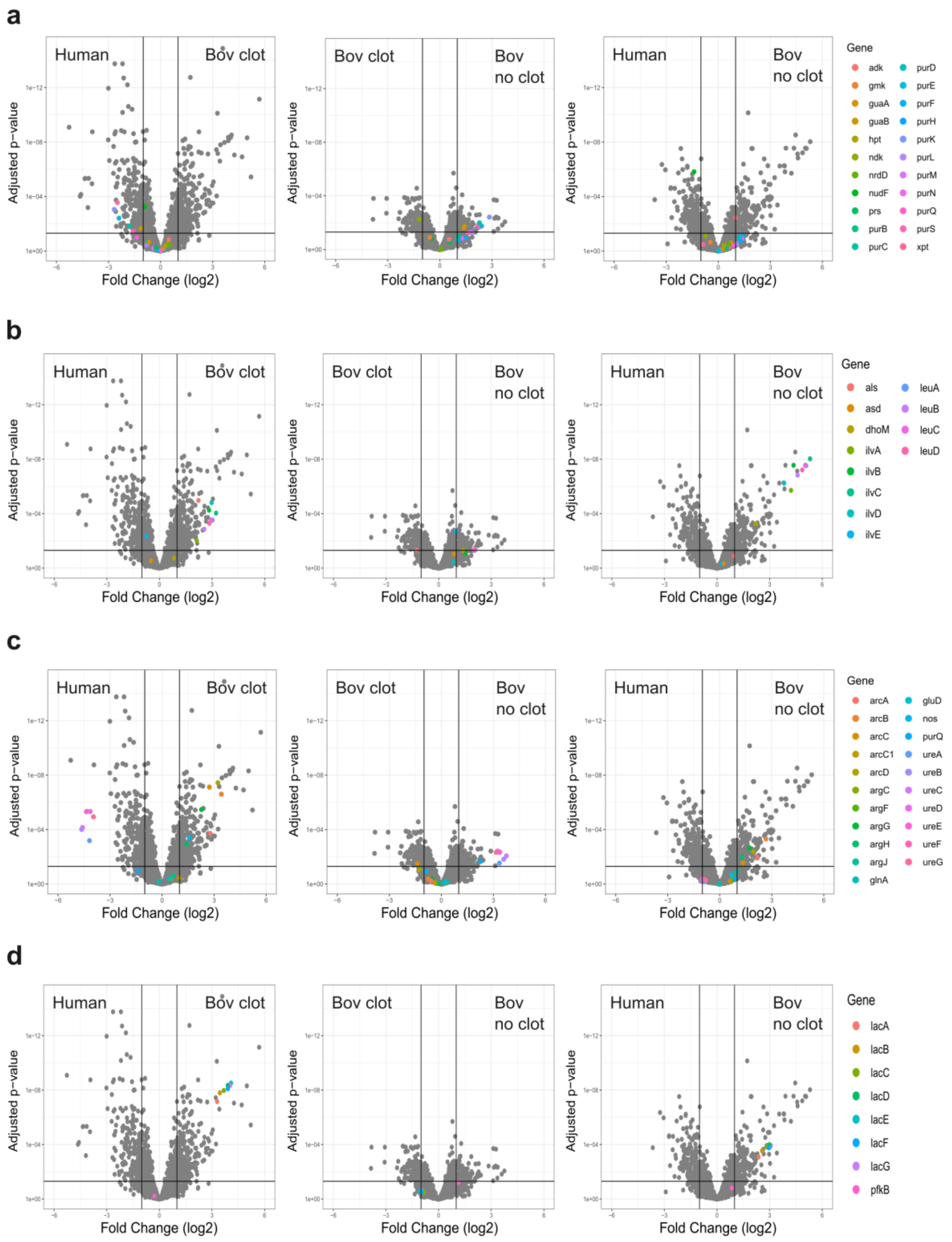


**Supp. Fig. 2| Volcano plots demonstrating the metabolic pathways differentially expressed between groups in milk. a,** Purine metabolism pathway. **b,** Valine, leucine, and isoleucine biosynthesis. **c,** Arginine biosynthesis. **d,** Galactose metabolism.

**a** **Purine Metabolism Pathway in RF122**

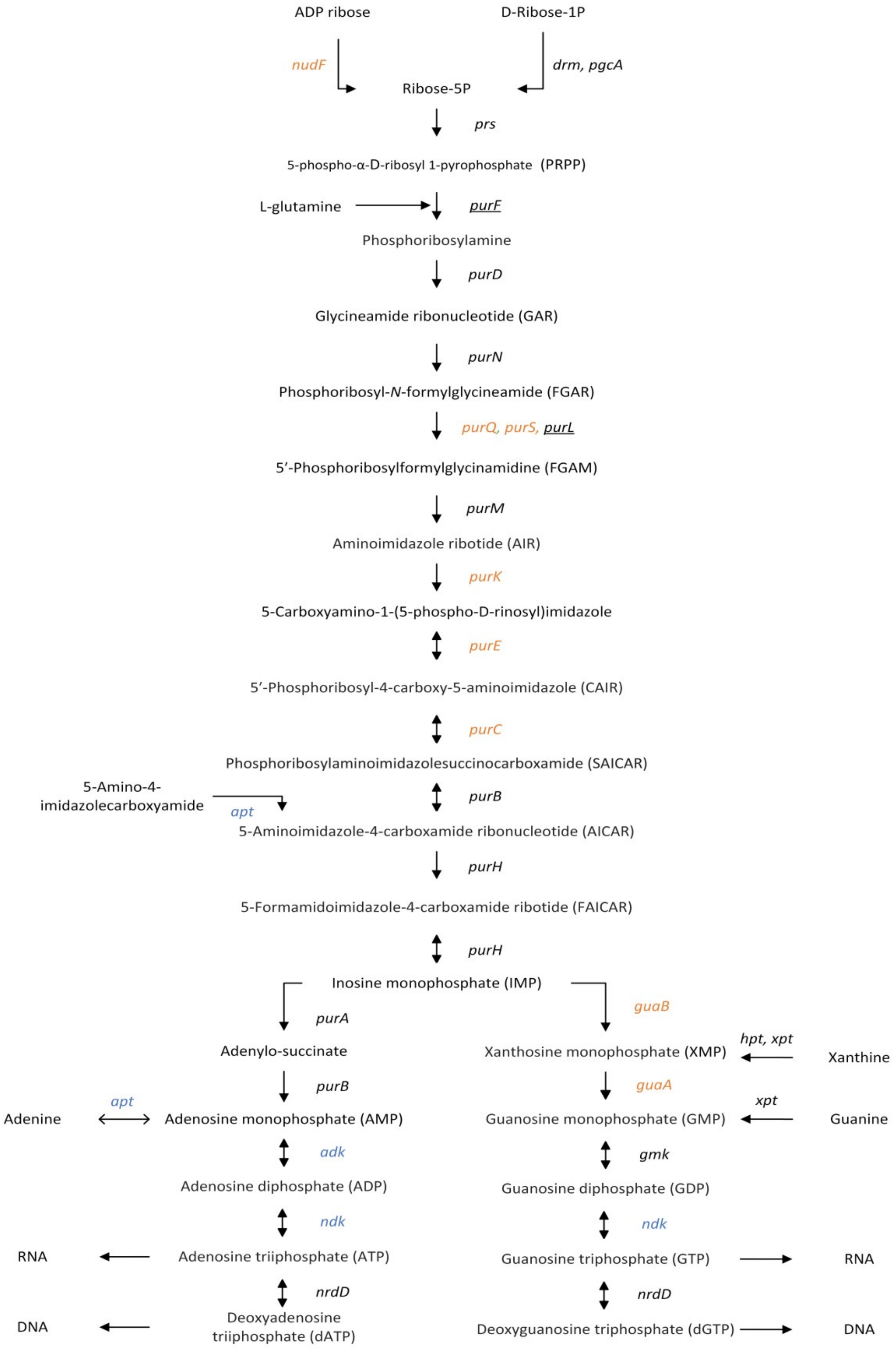

**b** **Valine, Leucine, and Isoleucine Biosynthesis Pathway in RF122**

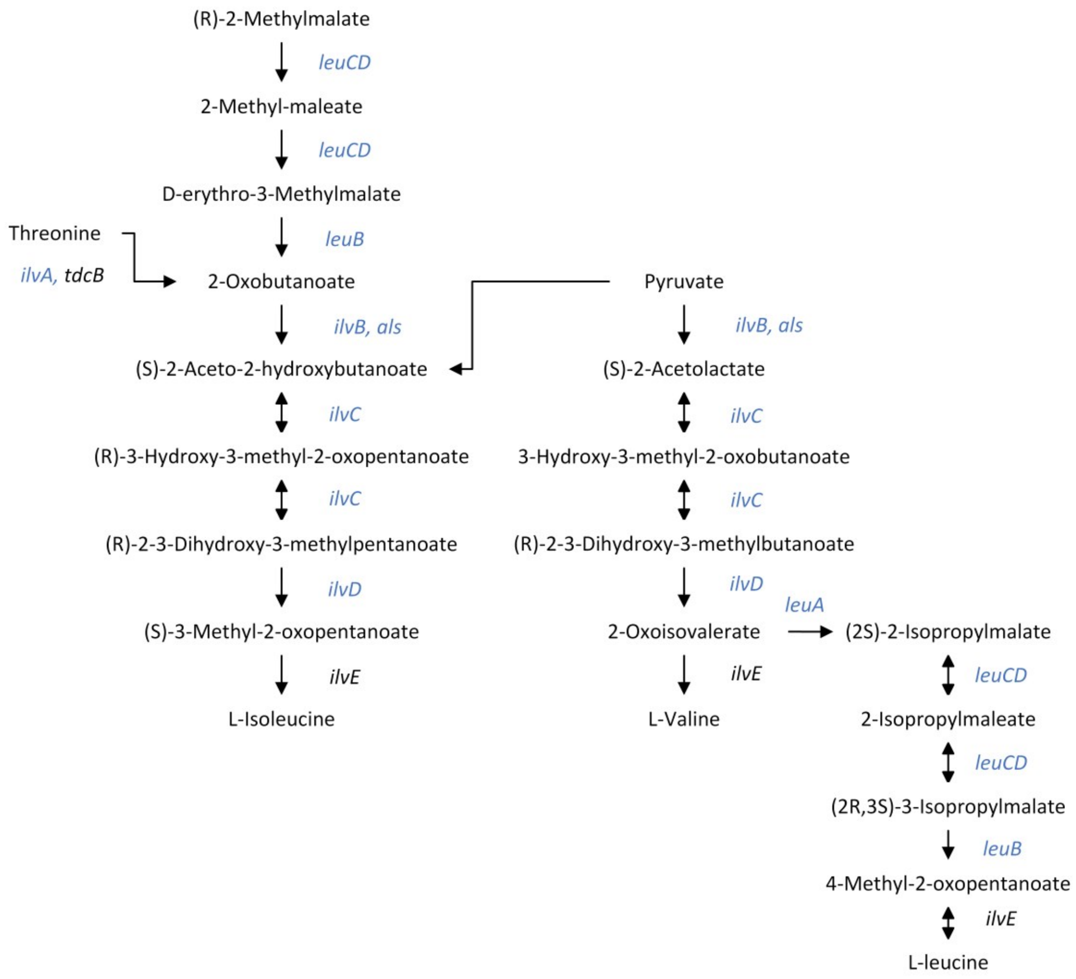


**c** **Arginine Biosynthesis Pathway in RF122**

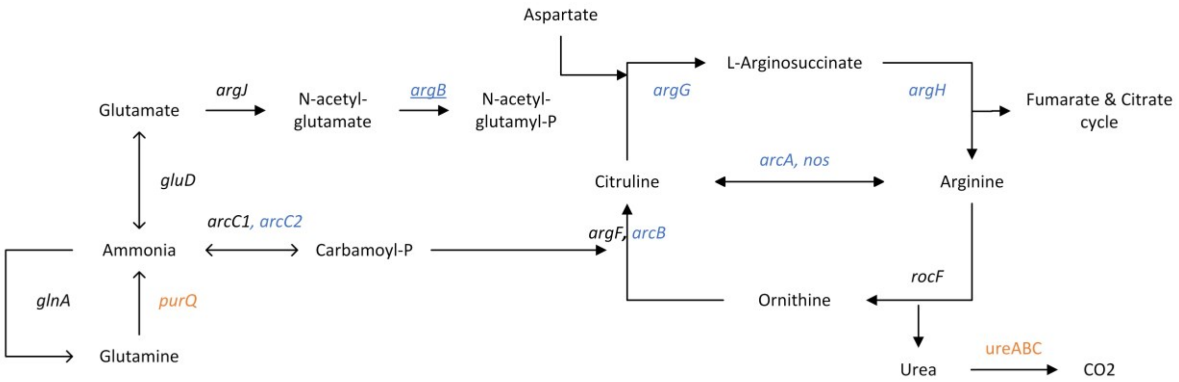

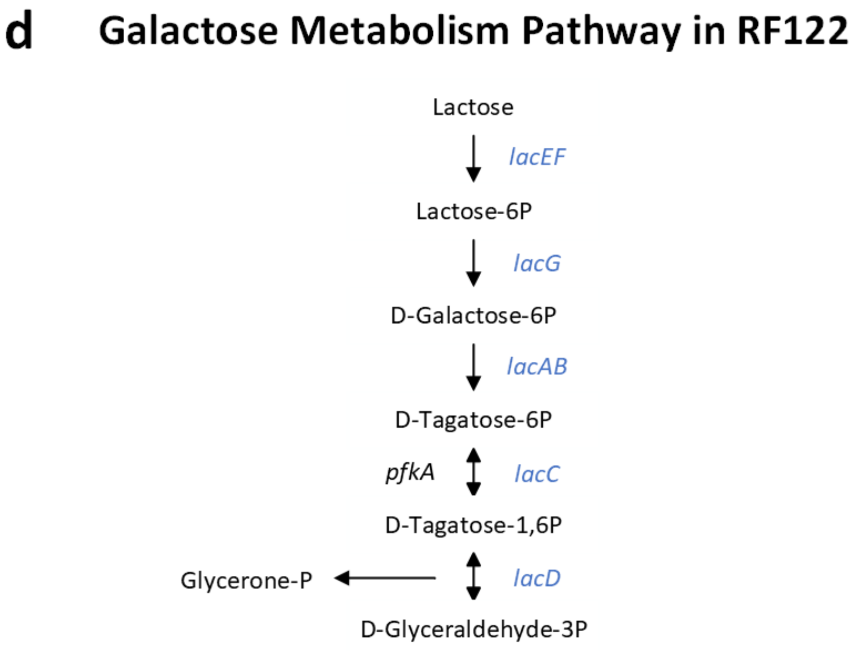


**Supp. Fig. 3| Schematics of the major metabolic pathways differentially expressed between groups in milk. a,** Purine metabolism pathway. **b,** Valine, leucine, and isoleucine biosynthesis. **c,** Arginine biosynthesis. **d,** Galactose metabolism. Genes coloured in orange are more highly expressed in human or non-clotting strains. Genes coloured in blue are more highly expressed in bovine strains compared to human strains. Genes that are underlined, were also identified to contain SNPs in the pairwise SNP analysis.

| Strain | WT | pALC2073 | pCT::*aur* | pCT::*sspA* |
|---|---|---|---|---|
| USA300 | 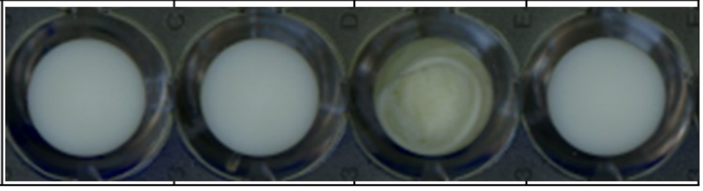 | | | |

**Supp. Fig. 4| Overexpression of aureolysin provides milk clotting of *S. aureus* USA300.** Arla Cravendale filtered whole milk was inoculated with each strain in triplicate and incubated statically at 37°C for 24 h. Milk clotting was assessed visually and imaged using an Epson scanner.

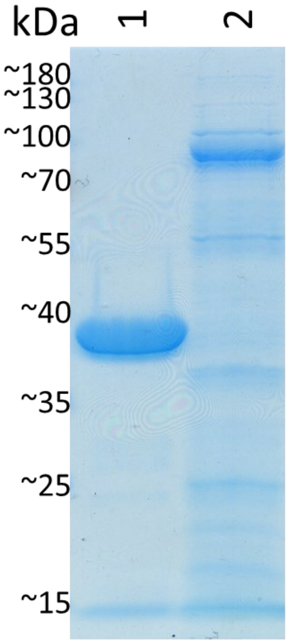


**Supp. Fig. 5| Concentrated supernatant of USA300Δprotease pCT::*aur*.** SDS-PAGE showing the protein profile of (1) concentrated supernatant of USA300Δprotease pCT::*aur* and (2) USA300Δprotease pALC2073.

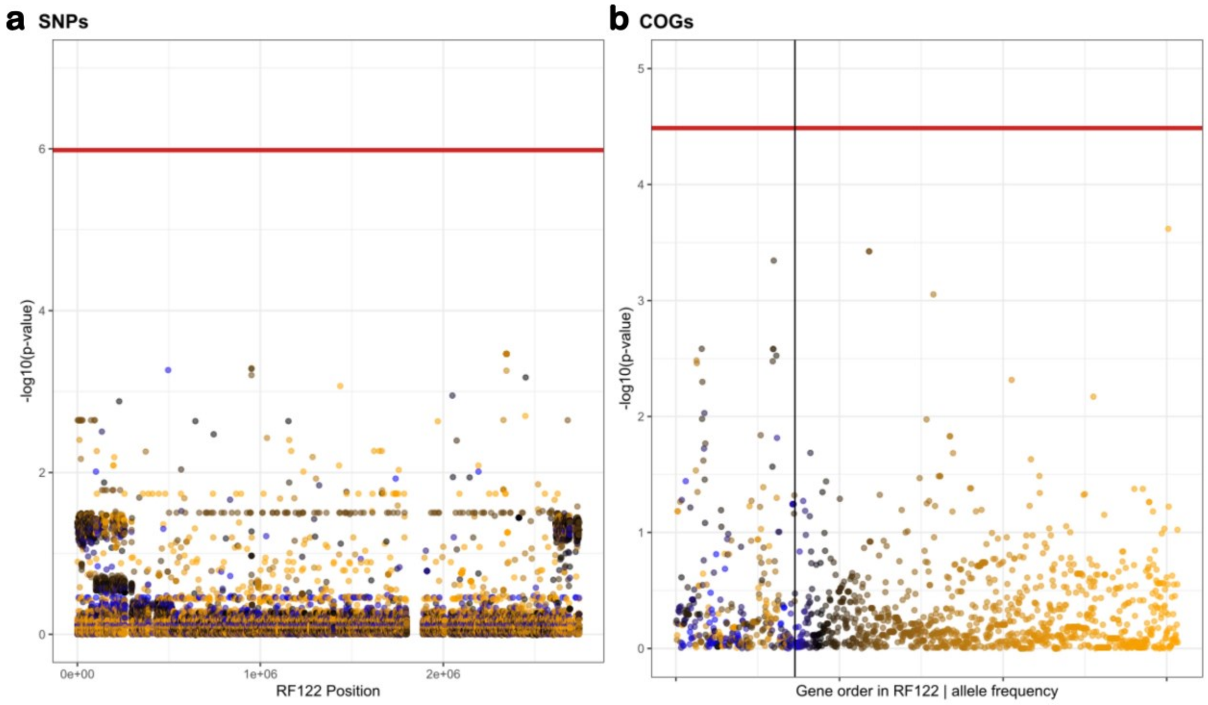


**Supp. Fig. 6| Genome wide association analysis does not identify a genetic basis for the milk clotting phenotype within lineages, suggesting convergent evolution. a,** Manhattan plot showing the significance of association of the core SNPs used to generate the phylogeny in Fig 1. The co-ordinates of SNPs in the reference genome (bp) are shown on the X axis. The Y axis shows the absolute magnitude of p-values for each associated variant. Each point represents a single variant and is coloured proportionally to its frequency across the data, as indicated by the scale bar. A horizontal red line shows the Bonferroni-corrected significance threshold used (alpha=0.05) **b,** Manhattan plot [as in **a)**] but showing the significance of association for different COGs, each represented by a single point. COGs that are present in RF122 are ordered by their synteny in RF122, whereas the COGs that are absent in RF122 (to the right-hand side of the black vertical line) are ordered by their frequency across the entire dataset from most to least common.
